# Disruption of maternal ACKR3 has profound effects on embryos and offspring

**DOI:** 10.1101/2023.06.15.545186

**Authors:** Ayumi Fukuoka, Gillian J. Wilson, Elise Pitmon, Lily Koumbas Foley, Hanna Johnsson, Marieke Pingen, Gerard J. Graham

**Affiliations:** Chemokine Research Group, School of Infection and Immunity, College of Medical, Veterinary and Life Sciences, University of Glasgow, 120 University Place, Glasgow G12 8TA, UK

**Keywords:** atypical chemokine receptor, hematopoietic stem cells, development, immune responses

## Abstract

ACKR3, scavenges and degrades the stem cell recruiting chemokine CXCL12 which is essential for proper embryonic, and in particular hematopoietic, development. Here we demonstrate strong expression of ACKR3 on trophoblasts. Using a pharmacological blocker we demonstrate that ACKR3 is essential for preventing movement of CXCL12 from the mother to the embryo with elevated plasma CXCL12 levels being detected in embryos from ACKR3-blocker treated mothers. Mice born to mothers treated with the blocker are lighter and shorter than those born to vehicle-treated mothers and, in addition, display profound anaemia associated with a markedly reduced bone marrow hematopoietic stem cell population. Importantly, whilst the hematopoietic abnormalities are corrected as mice age, our studies reveal a postnatal window during which offspring of ACKR3 blocker treated mice are unable to mount effective inflammatory responses to inflammatory/infectious stimuli. Overall, these data demonstrate that ACKR3 is essential for preventing CXCL12 transfer from mother to embryo and for ensuring properly regulated CXCL12 control over the development of the hematopoietic system.

## Introduction

The migration of leukocytes is controlled predominantly by a family of cytokines, called chemokines, and their receptors(Bachelerie *et al*, 2014a; Sokol & Luster, 2015). The chemokine family is defined on the basis of a conserved cysteine motif and divided into 4 sub-families according to the specific nature of this motif (CC, CXC, XC and CX3C families)(Zlotnik & Yoshie, 2000).

The chemokine family emerged in early vertebrates(Nomiyama *et al*, 2011; Zlotnik *et al*, 2006) and the primordial chemokine is believed to be CXCL12(Karpova & Bonig, 2015; Lewellis & Knaut, 2012; Zlotnik *et al*., 2006). Whilst mice with a homozygous-null deletion in either CXCL12, or its receptor CXCR4(Nagasawa *et al*, 1996; Tachibana *et al*, 1998; Zou *et al*, 1998) die, perinatally, analyses indicate that they have severely depleted bone marrow (BM) hematopoiesis as well as numerous other abnormalities including disrupted vascular development and alterations to cortical interneuron development(Stumm & Höllt, 2007). Further analysis using zebrafish has shown that primordial germ cells also express CXCR4 and that animals with homozygous-null deletion in CXCL12 or CXCR4 are effectively sterile(Doitsidou *et al*, 2002; Molyneaux *et al*, 2003). Thus, the pairing of CXCL12 and CXCR4 is essential for stem cell migration, and the development of numerous tissue systems within the embryo, and interfering with this axis has profound implications for offspring.

In addition to the classical chemokine receptors there exists a subfamily of receptors called atypical chemokine receptors (ACKRs)(Bachelerie *et al*, 2014b; Bonecchi & Graham, 2016; Nibbs & Graham, 2013). These are 7-transmembrane spanning receptors but lack the typical signalling capabilities of the other chemokine receptors. There are currently 4 members of this family, labelled ACKR1-4. Functionally, with the exception of ACKR1, all ACKRs actively internalise and scavenge their ligands through lysosomal-driven degradation(Bonecchi *et al*, 2004; Bryce *et al*, 2016; Naumann *et al*, 2010; Weber *et al*, 2004). They are therefore involved in removing chemokines in specific tissue contexts and in sculpting chemokine gradients. In terms of ligands, ACKR3 predominantly binds, internalises and scavenges the primordial chemokine CXCL12. ACKR3 knockout is associated with perinatal lethality linked to disrupted cardiac development(Gerrits *et al*, 2008; Sierro *et al*, 2007; Yu *et al*, 2011). ACKR3 therefore presents itself as an important regulator of the *in vivo* activities of the chemokine-receptor pairing of CXCL12 and CXCR4.

We have previously demonstrated that ACKR2 is expressed on trophoblasts in the junctionalzone and that its primary function here is to ensure that the mother can mount systemic chemokine-driven responses without these ligands entering the fetal circulation and interfering with cell migration within the embryo(Lee *et al*, 2019). We refer to this as ‘chemokine compartmentalisation’ and it involves trophoblast ACKR2 scavenging its ligands on the maternal face of the placenta thereby preventing their transplacental entry into the embryonic circulation.

Here we demonstrate that ACKR3 is expressed in syncytiotrophoblasts and is essential for compartmentalisation of CXCL12 activity and that pharmacological inhibition of maternal ACKR3 is associated with increased levels of maternally-derived CXCL12 in embryonic plasma. This impacts general embryonic development and severely blunts the seeding of BM by hematopoietic stem (HSC) and progenitor cells (HPC). Pups born to mothers exposed to ACKR3 blockade are smaller than those born to control mothers. Furthermore they display profound anaemia and hematopoietic insufficiency which is associated with an inability to mount protective inflammatory responses to bacterial stimuli. This study, therefore, demonstrates an essential role for placental ACKR3 in compartmentalising CXCL12 activity on the maternal side of the placenta thereby ensuring proper hematopoiesis and immune response in offspring.

## Results

### ACKR3 is expressed in placental trophoblasts

We used QPCR to examine ACKR3 expression in placenta and maternal decidua from wild-type (WT) pregnant mice at various times post conception. ACKR3 expression was detected in maternal decidua and not altered at all time points (Fig 1A). On the other hand, increased expression was apparent in the placenta starting at E12.5 and continuing at a similar level until E16.5. Using *Ackr3^GFP^* reporter mice, along with co-staining for syncytiotrophoblast markers, CD9 and MCT4, expression of ACKR3 was localised to the labyrinth region of the mouse placenta and specifically expressed on syncytiotrophoblasts, which form a barrier between maternal blood and embryonic tissues (Figs 1B and EV1A). Furthermore, flow cytometric analysis indicated clear association of ACKR3-positive placental cells with the syncytiotrophoblast markers, CD9 and CD71, but lack of association with CD45 (hematopoietic cells), CD31 (endothelial cells), CD90 (fibroblasts) and EpCAM (epithelial cells) (Figs 1C, EV1B and EV1C). In addition, immunostaining of human placenta localises ACKR3 expression to syncytiotrophoblasts (Figs 1D and EV2A) confirming that trophoblast expression of ACKR3 is evolutionarily maintained within mammals. Thus, ACKR3 is expressed predominantly in the placenta and its expression is restricted to syncytiotrophoblasts.

**Figure 1.**
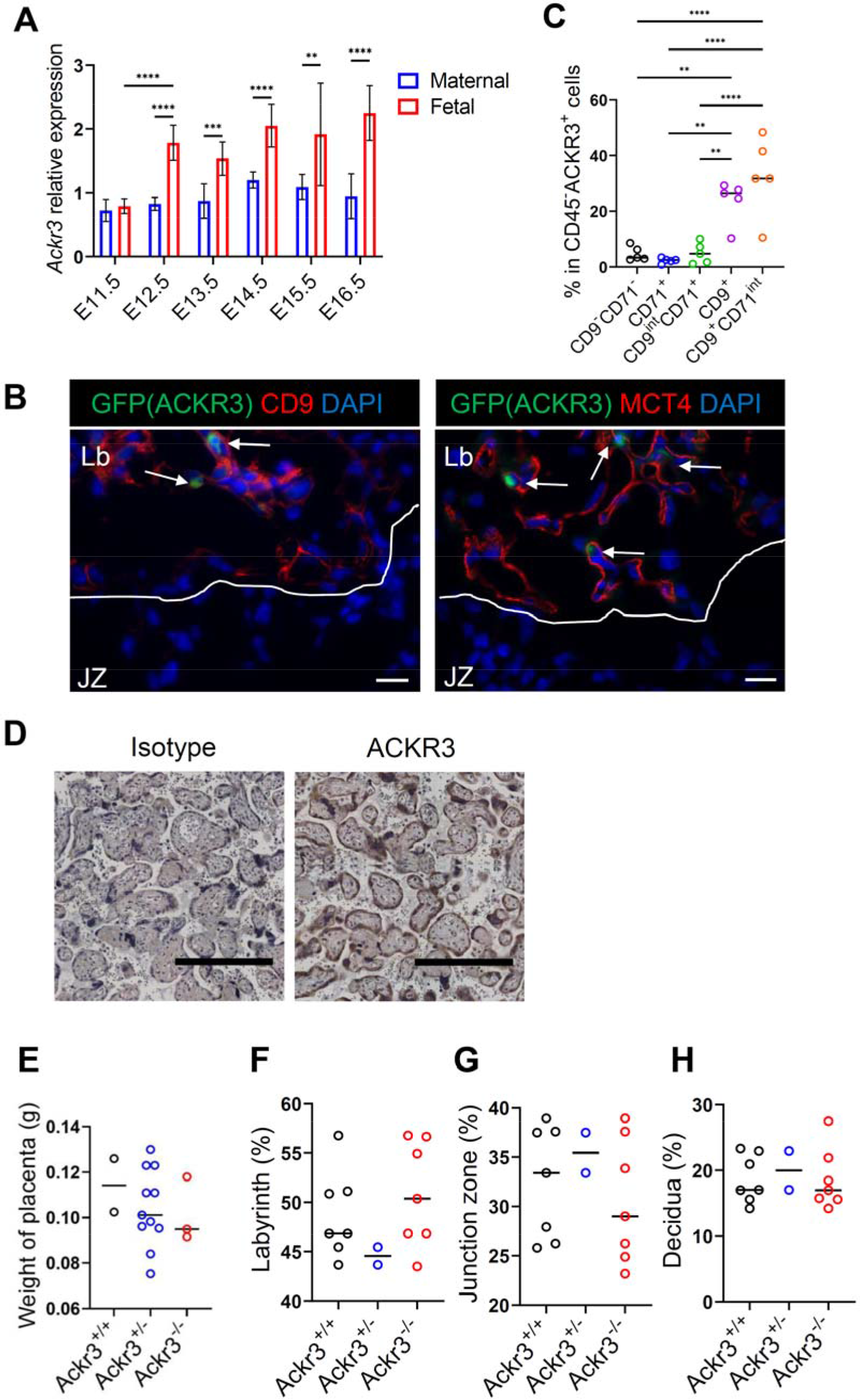
ACKR3 is expressed in the placenta. **A** Relative expression levels of mRNA of *Ackr3* in mouse placentas, and maternal decidua, between E11.5 and E16.5. **B** Representative images of placentas of ACKR3 reporter mice at E15.5. Sections were stained with syncytiotrophoblast (SynT) markers. Left image, Red:CD9 (SynT-I and -II), Blue:DAPI, Green:ACKR3, Right image, Red:Mct4 (SynT-II), Blue:DAPI, Green:ACKR3. Lb, Labyrinth, JZ, Junction zone. Scale bars indicate 20 μm. **C** Representative flow cytometry plots of trophoblast cells in ACKR3 reporter mice. The graph shows percentages of CD9^-^CD71^-^, CD71^+^, CD9^int^CD71^+^, CD9^+^, CD9^+^CD71^int^ cells in ACKR3^+^ cells in fetal side of placentas. **D** Representative images of immunohistochemistry of human placentas. Sections were stained with an anti-human ACKR3 antibody and counterstained with hematoxylin. Scale bars indicate 0.5 cm. **E** Weight of placentas at E15.5. **F-H** Histological analysis of placentas at E15.5. The graphs shows proportions of labyrinth (F), junction zone (G) and decidua (H). Data are representative of at least 2 independent experiments. Means ± SD with ****p < 0.001, ***p < 0.005, **p < 0.01, *p < 0.05 by Two-way ANOVA with Bonferroni’s post-test (A) or One-way ANOVA with Tukey’s post-test (C and E-H) are shown. See also Fig EV1 and 2

To investigate whether deletion of ACKR3 affects placental development or structure, we carried out gross analysis of placentas taken from WT, heterozygous (*Ackr3^GFP/+^*) and homozygous (*Ackr3^GFP/GFP^*) litter mates. This gross analysis did not reveal any significant differences in overall placental structure across the different genotypes (Fig EV2B). To analyse placental defects more precisely, we compared placental weight between the genotypes (Fig 1E) and the percentages of placental cross-section taken up by the decidua, labyrinth and junctional zones. None of these parameters where significantly different between the genotypes (Figs 1F, G and H). Therefore, the absence of ACKR3 appears to have no detectable gross impact on the placental development.

### ACKR3 regulates CXCL12 passage into the embryo

Next, we examined whether ACKR3 can compartmentalise chemokine function on the placenta. CXCL12 is detectable, at the transcript level, in both maternal decidua and placenta (Fig 2A). To determine roles for placental ACKR3 in limiting CXCL12 movement from maternal circulation to the embryo, we crossed heterozygous male and female mice (note that ACKR3 homozygosity is perinatally lethal(Gerrits *et al*., 2008; Sierro *et al*., 2007; Yu *et al*., 2011)) to give rise to WT, heterozygous and knockout embryos within each litter. We used an ELISA which is specific for murine CXCL12 to assess alterations in CXCL12 levels in plasma of *Ackr3*-deficient embryos.

**Figure 2.**
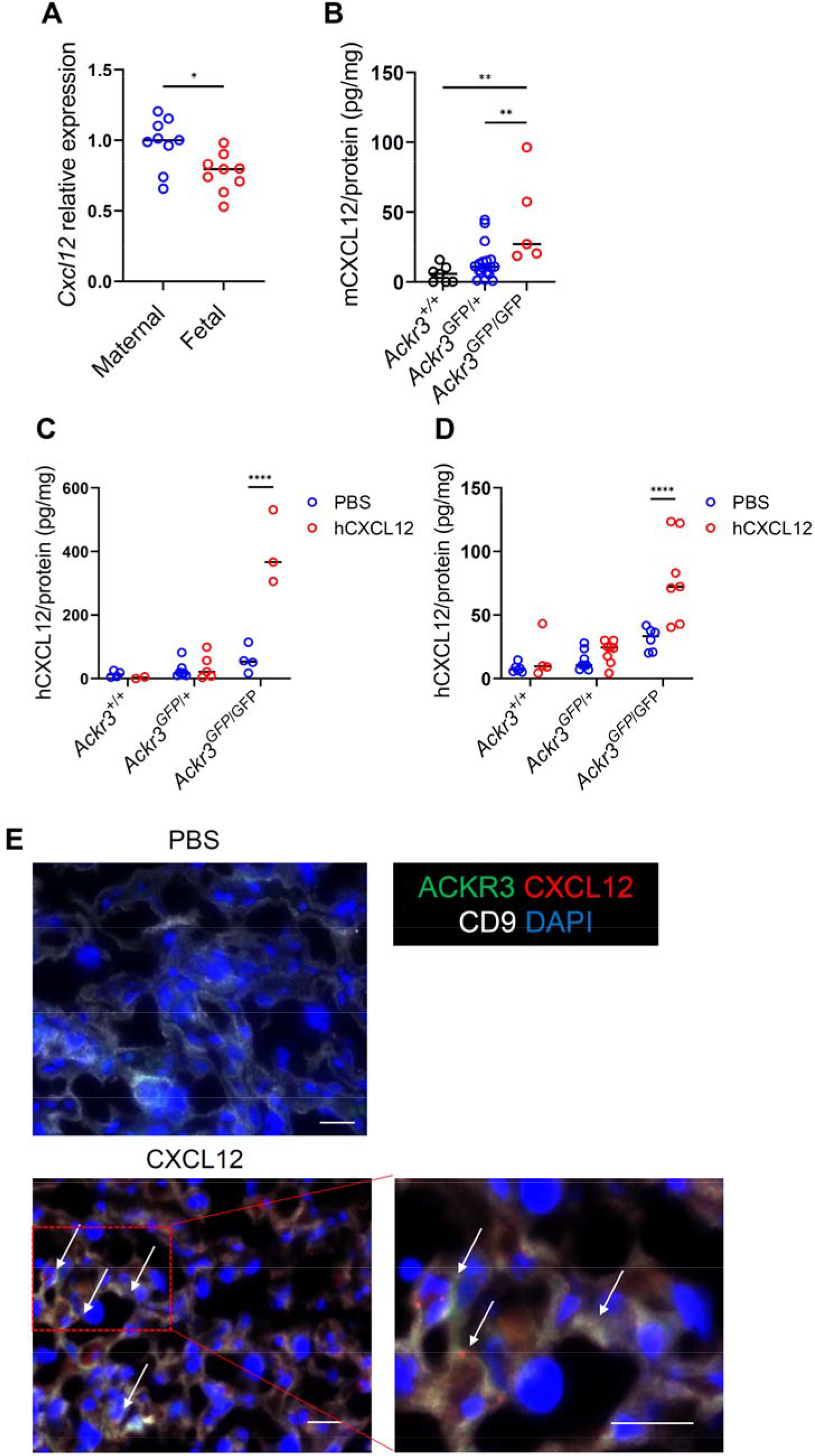
Placental ACKR3 prevents maternal CXCL12 entry into the embryonic circulation. **A** Relative expression levels of *Cxcl12* mRNA in maternal decidua (maternal) and placenta (fetal) side of isolated WT placentas at E15.5. **B** Mouse CXCL12 protein levels in plasma of embryos at E13.5 normalised to mgs of total protein. **C, D** CXCL12 levels, measured by ELISA, in plasma of embryos at E12.5 (C) and E15.5 (D) after hCXCL12 injection and normalised to mgs of protein. **E** *Ackr3*^GFP/+^ placentas were obtained after Alexa647-conjugated CXCL12 and sections were stained with anti-CD9 antibody. Green: ACKR3, White: CD9, Red: CXCL12, Blue: DAPI. Scale bars indicate 20 μm. Data are representative of at least 2 independent experiments (A, C and D) or were pooled from 2 independent experiments (B). Means ± SD with ****p < 0.001, *p < 0.05 by Student’s t test (A), One-way ANOVA with Tukey’s post-test (B) or Two-way ANOVA with Bonferroni’s post-test (C and D) are shown.

Plasma CXCL12 levels at E13.5 were significantly higher in *Ackr3*-deficient embryos compared to WT embryos (Fig 2B). These data indicate that ACKR3 regulates CXCL12 levels in embryos.

To examine the ability of ACKR3 to prevent CXCL12 passage from the mother to the embryo, we next injected *Ackr3*^GFP/+^ pregnant mothers crossed with *Ackr3*^GFP/+^ males with human CXCL12. The species-specific ELISA was used to measure chemokine movement from maternal plasma to fetal plasma. Human CXCL12 was readily detectable at both E12.5 and E15.5 in knockout but not WT embryonic plasma (Figs 2C and D).

Finally, to examine whether ACKR3-expressing trophoblasts take up CXCL12, *Ackr3*^GFP/+^ placentas were stained with a syncytiotrophoblast marker after injection of AF647-labelled CXCL12. AF647- CXCL12 was clearly seen to localize with ACKR3-expressing trophoblasts (Fig 2E).

These data demonstrate that ACKR3 on trophoblasts regulates CXCL12 passage from the placenta and the maternal circulation into the embryo.

### Pharmacological blockade of ACKR3 substantially compromises embryonic growth

As ACKR3 deletion is perinatally lethal, we opted to examine the developmental relevance of placental ACKR3 using a well-characterised pharmacological blocker, CCX771(Zabel *et al*, 2009) (Fig 3A). Injection of WT mothers with CCX771 did not significantly affect the total number of newborns within each litter but was associated with an increase in the numbers of dead newborns per litter (Figs 3B and C). In addition, we observed a number of pale-looking neonates particularly when CCX771 was co-administered with CXCL12 to the mother (Fig 3C). Embryos in CCX771-treated mothers showed high CXCL12 levels in blood compared to control embryos, indicating that maternal CXCL12 entered the embryonic circulation following inhibition of ACKR3 (Fig EV3A). Strikingly, CCX771 administration was also associated with a marked and significant reduction in weight of neonates, especially when combined with maternally administered CXCL12 (Fig 3D). Analysis of pups at 2 weeks of age showed that this reduction in body weight was also apparent at this time (Fig 3E) alongside a reduction in the length of the pups (Figs 3F and EV3B) and in the weight of the fat pads (Fig 3G). Analysis of mouse weight up to 7 weeks showed that, particularly for female mice, pups born to mothers receiving both CCX771 and CXCL12 never recovered the weight disadvantage that they were born with (Figs 3H and I).

**Figure 3.**
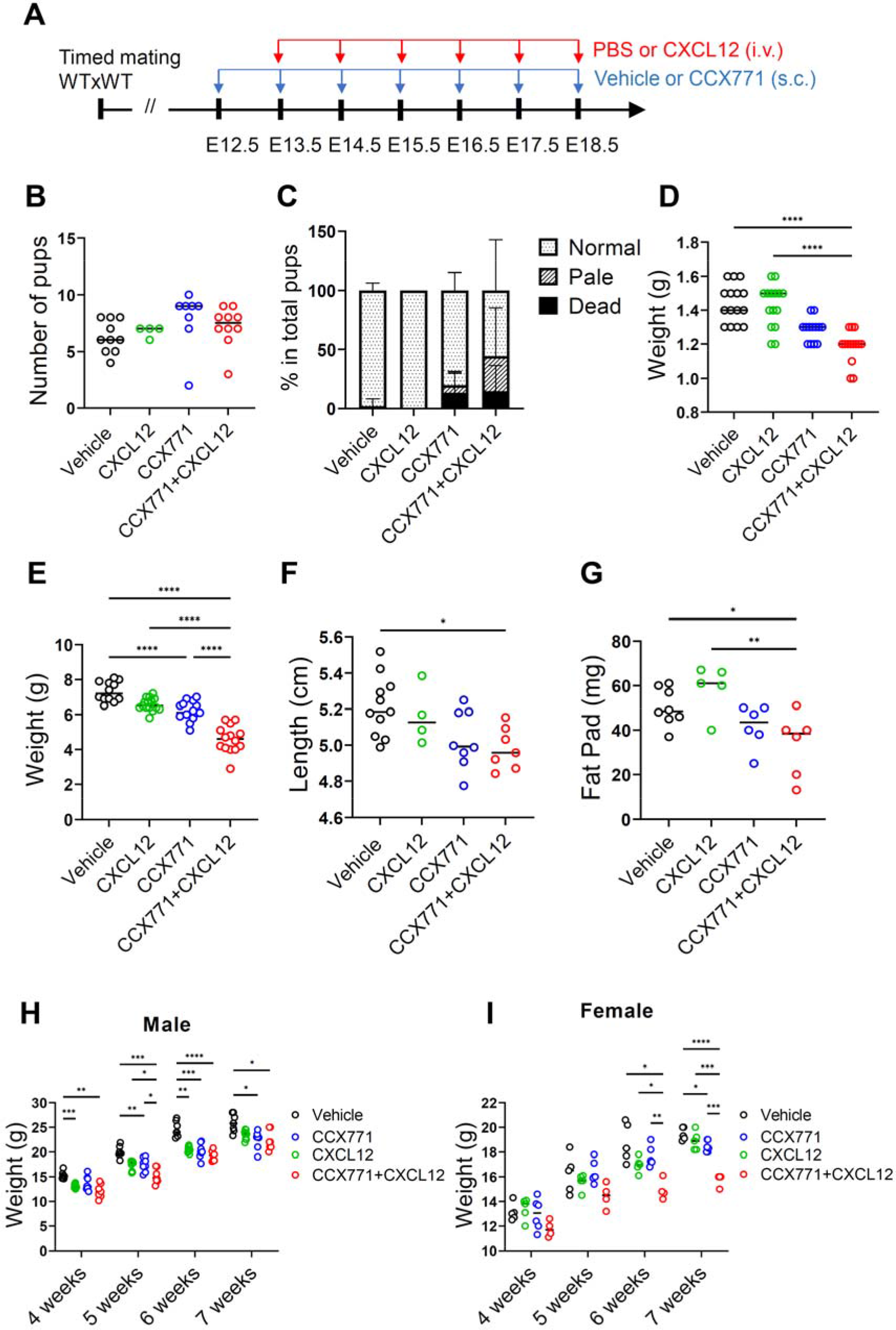
Blockade of ACKR3 during pregnancy affects embryonic development. **A** Experimental design. Wild-type (WT) pregnant mice were subcutaneously (s.c.) injected with Vehicle or CCX771 between E12.5 and E18.5. Additionally, the mice were intravenously (i.v.) injected with PBS or CXCL12 between E13.5 and E18.5. **B-D** Analysis of neonates born to the injected pregnant mice. Number of pups (B). The percentages of normal, pale and dead pups at postnatal day 0 (P0) (C). Weight of P0 pups (D). **E-G** Analysis of 2-week-old pups. Weight of 2-week-old pups (E). Length of pups (H). Weight of fat pads (G). **H,I** Weight of males (H) and females (I) between 4-week-old and 7-week-old. Data were pooled from 2 independent experiments. Means ± SD with ****p < 0.001, *p < 0.05 by One-way ANOVA with Tukey’s post-test (B, D and E-G) or Two-way ANOVA with Bonferroni’s post-test (H and I) are shown.

Importantly, mass spectrometry analysis of embryonic plasma revealed that CCX771 administered to the mother is frequently undetectable in the embryo and, where levels are detected, they were well below the IC_50_ for this blocker (18ng/ml). Additionally, where low level CCX771 was detected in embryos this did not positively correlate with decreased HSC population described below (Fig EV3C). This demonstrates that CCX771 does not cross the placenta in biologically significant levels indicating that its ACKR3 blocking activity is restricted to the mother and most likely to trophoblastic cells.

Overall, these data demonstrate that ACKR3 blockade, particularly in combination with maternally administered CXCL12, leads to embryonic compromise associated with reduced body weight which is maintained until maturity.

### ACKR3 blockade substantially impairs hematopoietic development

As mentioned above, neonates born to mothers injected with CCX771 or CCX771 and CXCL12 were observed to be paler than control neonates. Hematology analysis of neonates’ blood revealed a substantial reduction in red blood cell numbers in neonates born to mothers injected with CCX771 and CXCL12 (Fig 4A). Notably and in contrast to the data relating to weight and length described above, treatment of mothers with ACKR3 blocker alone led to similar levels of anaemia compared to those seen with a combination of blocker and CXCL12 (Fig 4B). In addition, a general reduction in white blood cell numbers was also seen and again this was seen for neonates from both blocker and blocker plus CXCL12 treated mothers (Fig 4C). Thus, the pale appearance of embryos from mothers injected with CCX771, and CCX771 plus CXCL12, is associated with anaemia.

**Figure 4.**
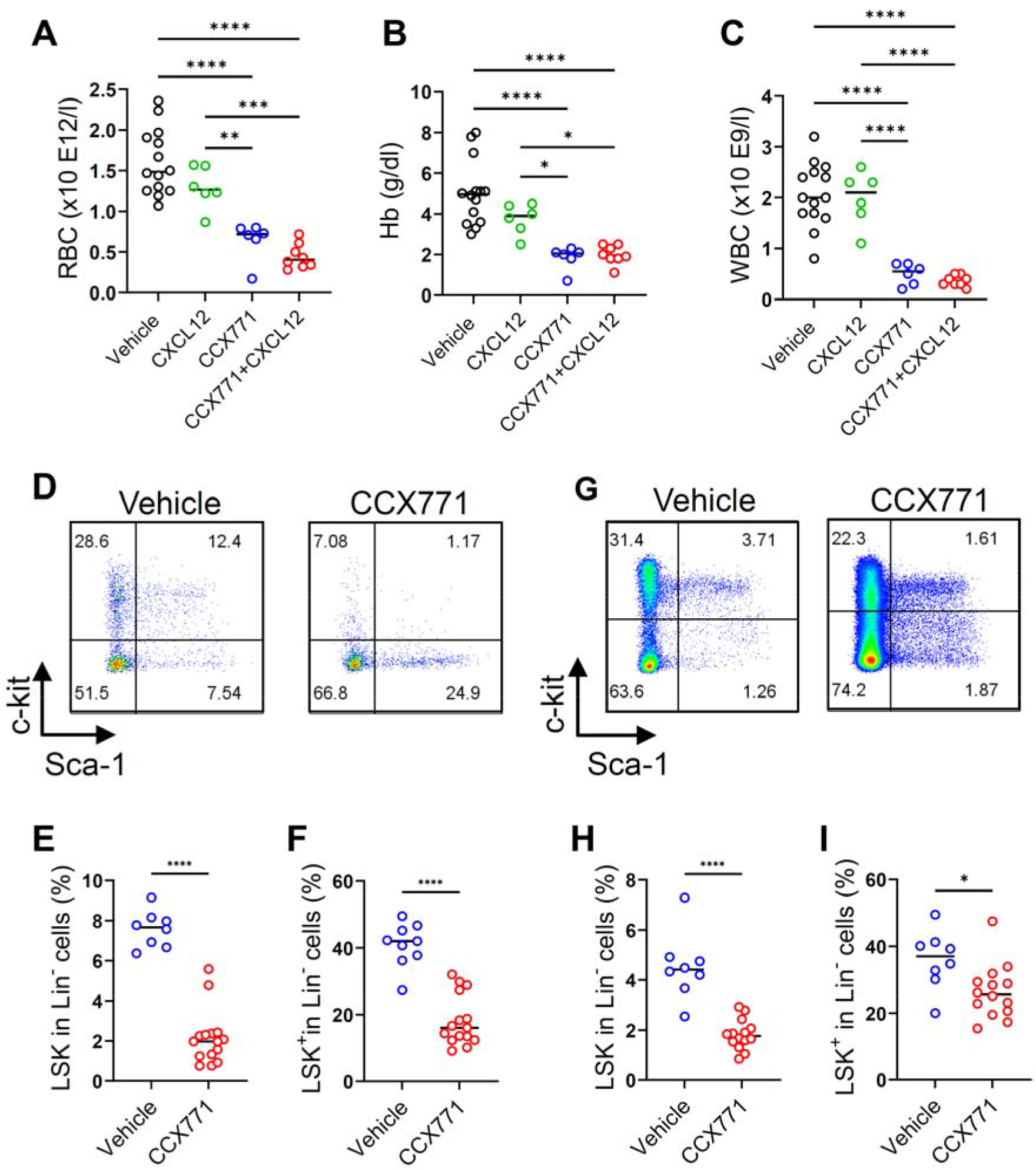
ACKR3 Blockade impairs HSC development in embryos. **A-C** Hematology blood tests of neonates born to mice injected with vehicle, CXCL12, CCX771 or CCX771 and CXCL12. Red blood cells (RBC) (A), haemoglobin (Hb) (B) and white blood cells (WBC) (C). **D-I** Flow cytometry analysis of bone marrow (BM) and fetal liver (FL) of E18.5 embryos from vehicle or CCX771 injected pregnant mice. Representative flow cytometry plots of Lineage marker (Lin) negative cells in BM. The percentage of Lin^-^, Sca-1^+^ and c-kit^+^ (LSK) cells (E) and Lin^-^, Sca-1^-^ and c-kit^+^ (LSK^+^) cells (F) in BM. Representative flow cytometry plots of Lin^-^ cells in FL in E18.5 embryos (G). The percentage of LSK cells (H) and LSK^+^ cells (I) in FL. Data were pooled from 2 independent experiments. Means ± SD with ****p < 0.001, ***p < 0.005, **p < 0.01, *p < 0.05 by One-way ANOVA with Tukey’s post-test (A-C) or Student’s t test (E,F,H and I-J) are shown. See also Fig EV3.

To determine the basis for this phenotype we next examined BM and FL HSC numbers in embryos. This analysis focused on vehicle and CCX771-treated mothers as the hematopoietic anomalies described above were evident in mice treated only with CCX771. In BM, there was a highly significant reduction in the number of Lineage marker negative (Lin)^-^Sca-1^+^c-Kit^+^ (LSK) HSCs in the embryos from CCX771 treated, compared to vehicle treated mothers (Figs 4D and E). This was also associated with a decrease in the number of Lin^-^Sca-1^-^c-Kit^+^ (LS-K^+^) hematopoietic progenitor cells (HPC) within the BM (Fig 4F). Intriguingly, HSC and HPC populations were also reduced within the FL in embryos from CCX771-treated mothers (Figs 4G, H and I).

Next, we examined BM hematopoietic parameters in mice at various time points post birth. Neonates born to both CCX771 and CCX771 plus CXCL12 injected mothers displayed significant reduction in both HSC and HPC numbers (Figs 5A and B). At 2 weeks, pups born to CCX771-injected mothers had recovered numbers of HSC but remained severely depleted in HPC (Figs 5C and D). In contrast, both HSC and HPC numbers remained depleted in pups born to mothers treated with a combination of CCX771 and CXCL12. By 7 weeks, HSC and HPC numbers had normalised across progeny from all the treated groups (Fig 5E).

**Figure 5.**
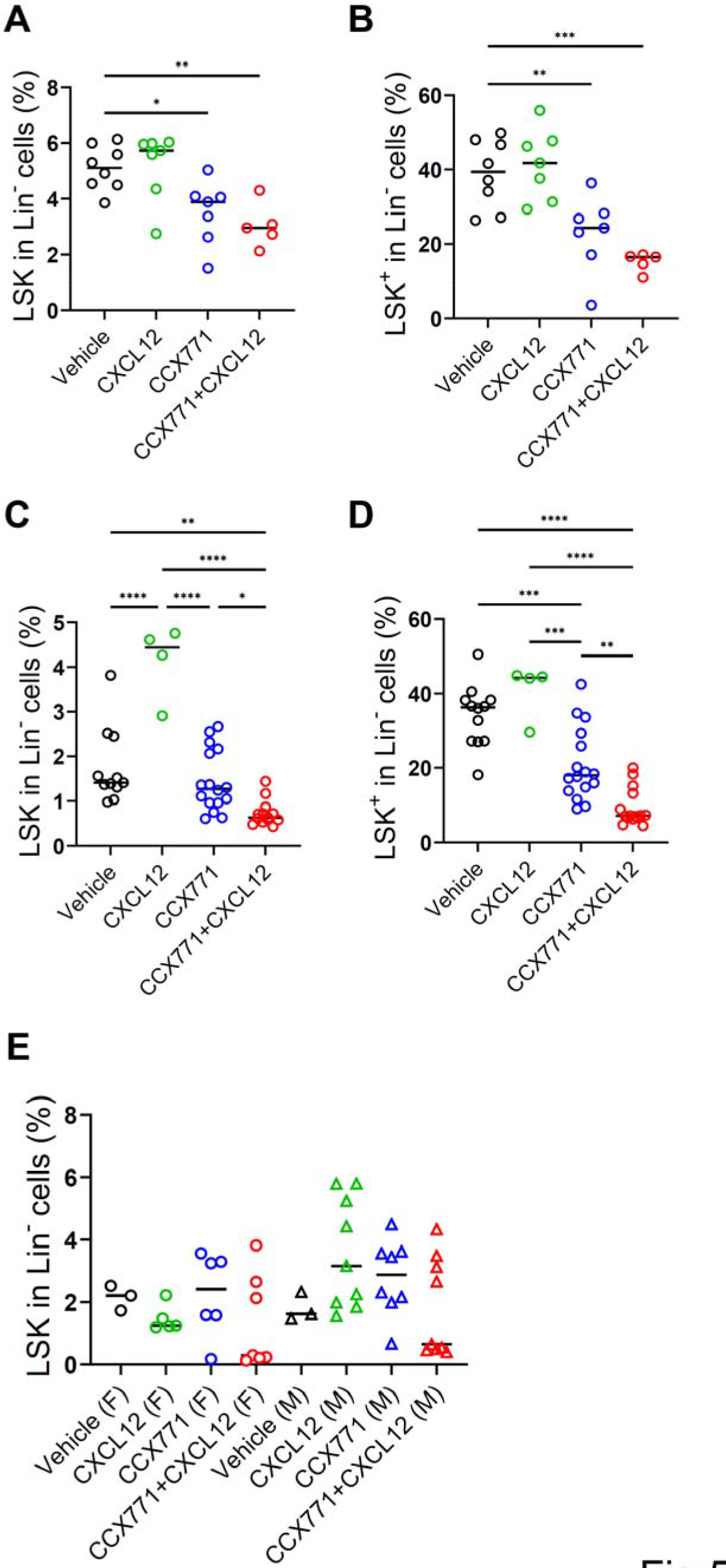
Maternal ACKR3 blockade is associated with sustained disrupted hematopoiesis after birth. Wild-type pregnant mice were injected with vehicle, CXCL12, CCX771 or CCX771 and CXCL12 (Figure 3A). **A, B** Analysis of hematopoietic stem cells in neonates. The percentage of lineage marker (Lin)^-^, Sca- 1^+^ and c-kit^+^ cells (LSK) (A) and Lin^-^, Sca-1^-^ and c-kit^+^ cells (LSK^+^) (B) in bone marrow (BM) in neonates. **C, D** The percentage of BM LSK cells (C) and LSK^+^ cells (D) in Lin^-^ cell in 2-week-old pups. **E** The percentage of BM LSK cells in Lin^-^ cells in 7-week-old mice. F, Females, M, Males. Data were pooled from 2 independent experiments. Means ± SD with ****p < 0.001, ***p < 0.005, **p < 0.01, *p < 0.05 by One-way ANOVA with Tukey’s post-test are shown.

These data demonstrate that ACKR3 blockade is associated with severely compromised hematopoietic development in embryos.

### ACKR3 blockade also has a significant impact on B cell numbers

Given known roles for CXCL12 in B cell biology(D’Apuzzo *et al*, 1997; Nagasawa *et al*., 1996) we analysed B cell numbers in offspring. In neonates, there was a general reduction in total B cell numbers, as well as numbers of BM IgM^+^ immature B cells in pups born from CCX771 and CCX771 plus CXCL12 treated mothers (Figs 6A-C). Reduction in numbers of B220^+^IgM^-^ B cells (Pro/Pre-B cells) in BM was seen only for pups born to mothers injected with a combination of CCX771 and CXCL12 (Fig 6D). At 2 weeks this reduction was seen for total B cell numbers, and for mature, but not immature B cells which had rebounded in number (Figs 6E-G and EV4). Both maternal CCX771 and CCX771 plus CXCL12 administration led to significant reduction in mature B cells in 2-week-old offspring (Fig 6F). By 7 weeks no differences in B-cell numbers were detected in any of the progeny from any of the treated groups (Figs 6H and I). These results suggest that maternal ACKR3 blockade also has an impact on early B cell development within offspring.

**Figure 6.**
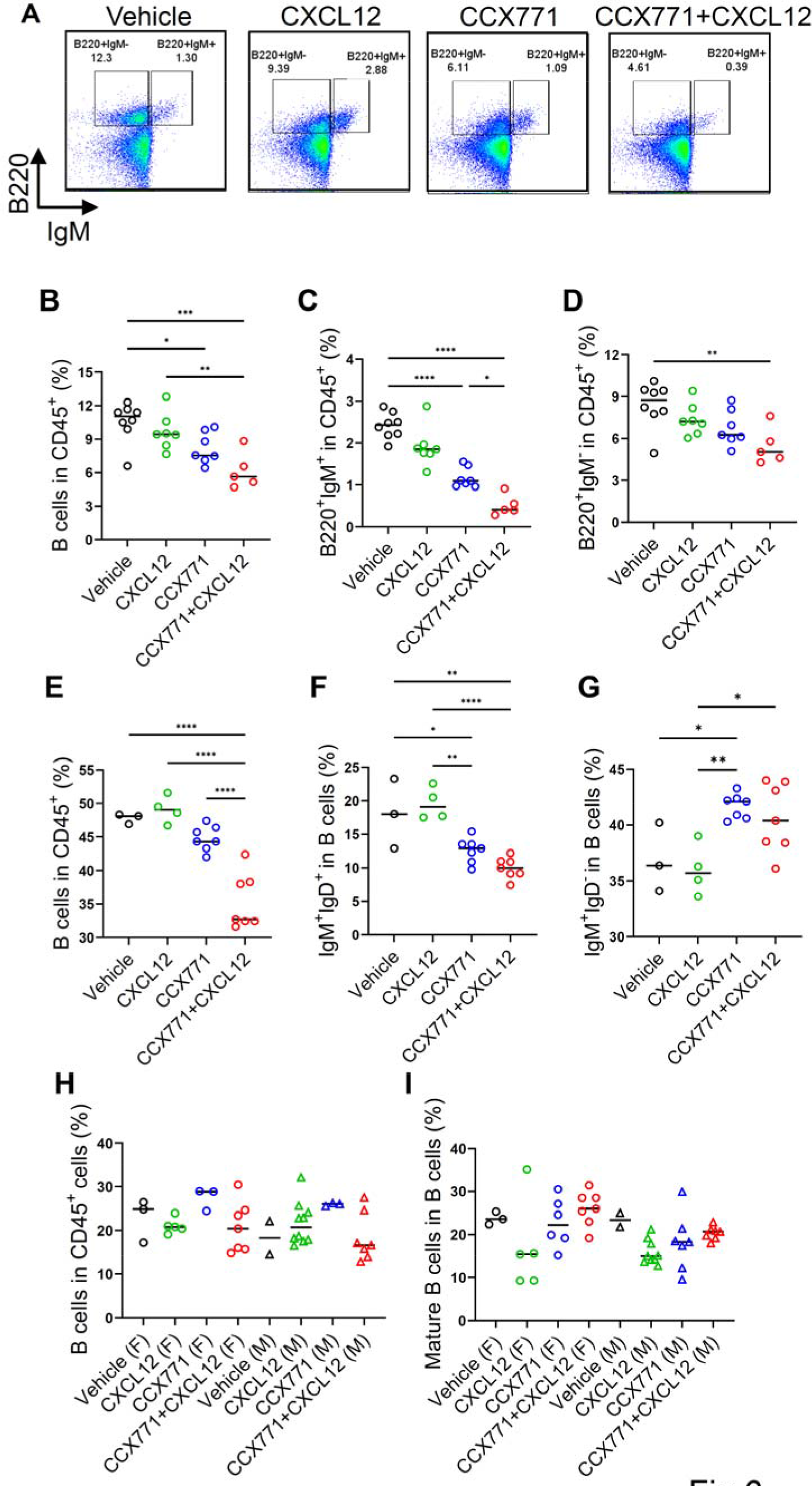
Maternal ACKR3 blockade is associated with decreased mature B cell numbers in offspring. **A-D** B cell populations in bone marrow (BM) in neonates born to wild-type pregnant mice injected with vehicle, CXCL12, CCX771 or CCX771 and CXCL12. Representative flow cytometry plots (A). The percentage of total B cells (B), immature/mature (B220^+^IgM^+^) cells (C) and Pro/Pre-B cells (B220^+^gM^-^) cells (D). **E-G** The percentage of total B cells (E), mature B cells (IgM^+^IgD^+^) cells (F) and immature B cells (IgM^+^IgD^-^) cells (G) in 2-week-old mice. **H, I** The percentage of total B cells (H) and mature cells (I) in 7-week-old offspring. F, females, M, males. Data are representative of 2 independent experiments (A-G). Data were pooled from 2 independent experiments (H and I). Means ± SD with ****p < 0.001, ***p < 0.005, **p < 0.01, *p < 0.05 by One-way ANOVA with Tukey’s post-test are shown. See also Fig EV4.

### ACKR3 blockade leads to deficiencies in the inflammatory response in neonates

Whilst the suppression of hematopoiesis seen in embryos and neonates from mothers treated with CCX771 resolved by 7 weeks, this leaves a critical postnatal period in which mice born to mothers treated with CCX771 have significantly reduced numbers of cells from across the hematopoietic spectrum. To examine the implications of this in terms of the ability of preweaned pups to mount protective inflammatory responses we challenged 2-week-old mice, born from mothers injected either with vehicle or CCX771, by intra-peritoneal administration of pHrodo Red conjugated *E. coli* particles. Pups born to vehicle-injected mothers displayed increased numbers of BM HSC and reduced HPC with evidence that these cells are mobilised from the BM in response to *E. coli* particle administration. These cell populations in pups born to mothers injected with CCX771 were substantially reduced although not altered by *E. coli* particles (Figs 7A and B). Ly6Chigh monocytes and neutrophils in the BM in pups born to CCX771-treated mothers were significantly lower than those in control pups. Mobilisation of Ly6Chigh monocytes by *E. coli* particle administration was detected in control pups, but not in pups born to CCX771-treated mothers, while mobilisation of neutrophils was observed in both groups. (Figs 7C and D).

**Figure 7.**
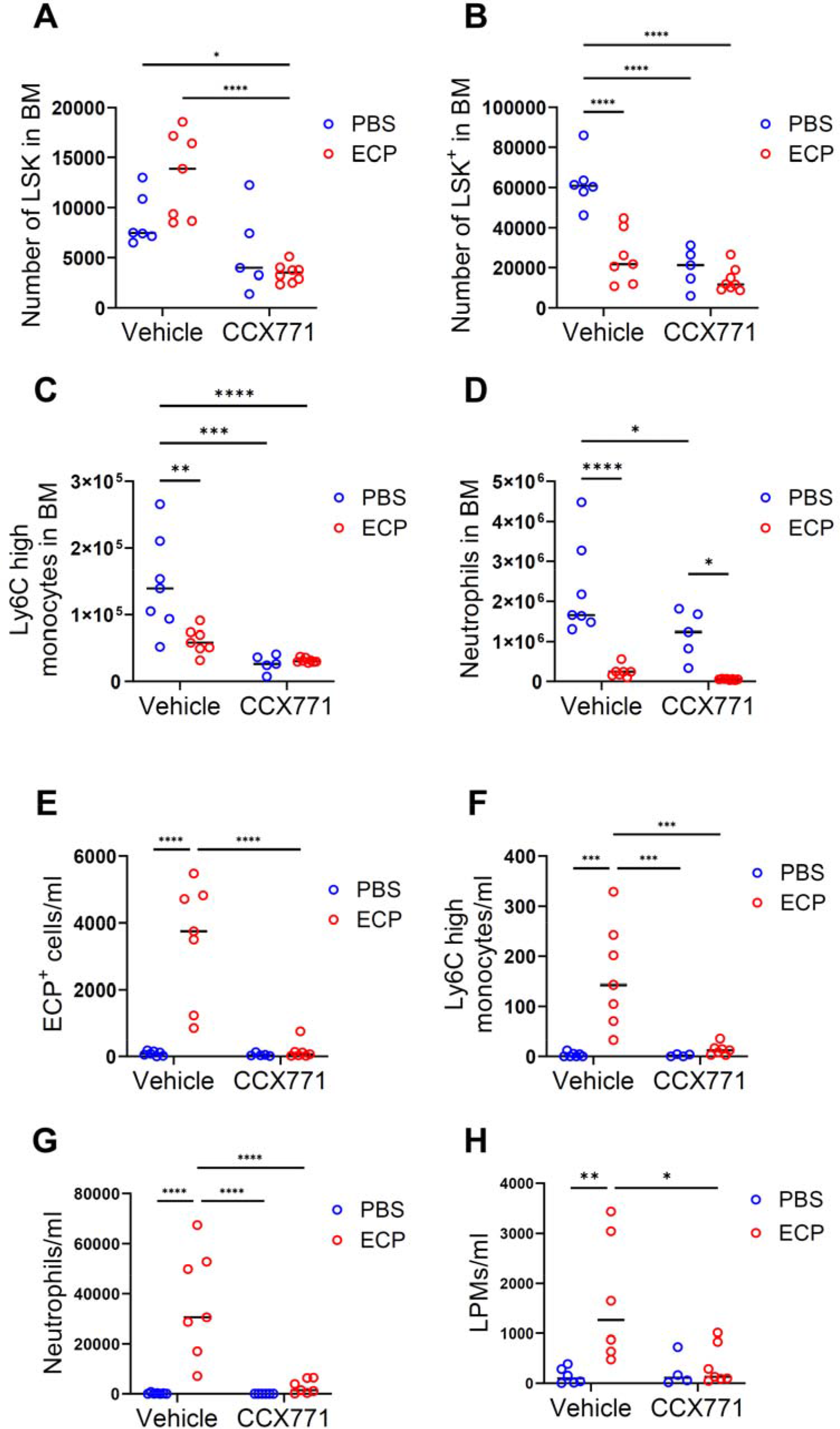
Maternal ACKR3 blockade is associated with impaired inflammatory responses in offspring. **A-H** 2-week-old pups born to wild-type pregnant mice injected with vehicle or CCX771 were intraperitoneally injected with PBS or pHrodo Red dye conjugated *E.coli* particles (ECP). 24 hours after the injection, the peritoneal lavage and the bone marrow (BM) cells were analyzed by flow cytometry. Total numbers of LSK (A) and LSK^+^ cells (B), Ly6C high monocytes (C) and neutrophils (D) in BM, and ECP^+^ cells (E), Ly6C high monocytes (F), neutrophils (G) and Large peritoneal macrophages (LPMs) (H) in peritoneal lavage. Data are representative of 2 independent experiments. Means ± SD with ****p < 0.001, ***p < 0.005, *p < 0.05 by Student’s t test are shown. ns indicates not significant. See also Fig EV5.

Peritoneal contents were collected by lavage, and *E. coli* particle uptake and recruited cells assessed by flow cytometry (Figs EV5A-E). Whilst there was substantial uptake of *E. coli* particles by recruited cells in the peritoneum of pups born to control mothers, this was severely depleted in the peritoneum of pups born to CCX771-treated mothers (Fig 7E). When recruited cells were analyzed it was seen that pups born to control treated mothers demonstrated significant recruitment of Ly6Chigh monocytes and neutrophils but that there was minimal recruitment of these cells in pups born to mothers treated with CCX771 (Figs 7F and G). These results suggest a fundamental inability to mount inflammatory responses in these offspring. Tissue-resident large peritoneal macrophages (LPMs) were increased in control pups by the administration of the *E. coli* particles, however, LPMs in pups born to CCX771-treated mothers were not altered (Figs 7H). In keeping with the data described above, there was a reduction in peritoneal B cell numbers in pups born to CCX771-treated mothers but these cells appear to be unaffected by the presence or absence of *E. coli* particles (Fig EV5F).

Overall, these data indicate that the depletion of hematopoietic stem and progenitor cells seen in neonates from CCX771-treated mothers is associated with a profound inability to recruit inflammatory cells in response to peritoneal bacterial challenge.

## Discussion

Here we demonstrate that ACKR3 has an important role in restricting the movement of CXCL12 across the trophoblast barrier into the embryonic circulation. It is clear from our analyses that the effects of interfering with ACKR3 are profound in terms of altering developmental parameters, with pups born to ACKR3 blocker treated mothers being characterised by growth retardation and marked alterations to hematopoietic system development. These phenotypes continue through the neonatal period and, whilst hematopoietic parameters are normalised by 7 weeks, the reduced weight seen in mice born to ACKR3 blocker treated mothers is not fully recovered over time. The normalisation of hematopoietic parameters is fully in keeping with the self-renewal potential of the hematopoietic system.

Notably, administration of CCX771 alone to mothers resulted in a significant reduction of HSC and HPC numbers and associated severe anaemia, whilst administration of a combination of CCX771 and CXCL12 to mothers was required for significant effects on more general developmental parameters including weight and length. This suggests that HSC migration within the embryo is more susceptible to low level maternal CXCL12 entry into the embryo than other developmental processes.

Crucially this means that there is a significant postnatal period during which mice born to mothers with impaired ACKR3 activity have significant deficiencies within their hematopoietic system. Importantly this is reflected in a marked inability of these mice to mount protective inflammatory responses to peritoneal bacterial challenge. This suggests that altered maternal ACKR3 activity would leave offspring susceptible to potentially lethal postnatal infections of the peritoneal cavity and other tissues.

It is worth highlighting that this study could not be done with fully ACKR3-deficient mothers as homozygous ACKR3 deficiency is perinatally lethal. Thus, the availability of a high-quality ACKR3 antagonist, which does not significantly cross the placenta has afforded us a unique opportunity to analyse chemokine compartmentalisation in the context of ACKR3.

Interestingly, whilst we have demonstrated a similar role for ACKR2 in the placenta, it is important to note that the localization of ACKR2 and ACKR3 in the placenta is different. ACKR3 is expressed in trophoblasts in the labyrinth, while ACKR2 expression is observed predominantly on trophoblasts in the junctional zone(Lee *et al*., 2019). Previous studies have demonstrated that CXCL12 is expressed in trophoblasts(Ren *et al*, 2012; Wu *et al*, 2004) rather than the maternal decidua, while CCL2, which is one of the ligands for ACKR2, is expressed in maternal decidua(Lin *et al*, 2022) and also seen at high levels in the blood of inflamed or infected mothers. This suggests that ACKR2 mainly scavenges its ligand arising in the placenta from maternal blood or the maternal decidua. In contrast, ACKR3 is required to prevent CXCL12 derived from intra-placental sources including trophoblasts, as well as from maternal blood, from infiltrating into the embryonic circulation. Thus, whilst operating in a mechanistically similar manner, ACKR2 and ACKR3 display discrete domains of expression, and therefore function, within the placenta which reflect the expression patterns of the ligands.

Overall, therefore, our results indicate a crucial role for ACKR3 in protecting the embryo from maternal CXCL12 and that interference with ACKR3 function leads to severe developmental/hematopoietic consequences for offspring. These results suggest that defects in trophoblast ACKR3 in humans, associated with impaired expression levels or function, may lead to developmental/hematopoietic abnormalities similar to those observed in the present study. Notably, previous studies have shown that decreased ACKR3 expression in placentas and increased CXCL12 levels in bloods were observed in pregnant women diagnosed with preeclampsia(Lu *et al*, 2016; Schanz *et al*, 2011). Also, it is likely that this will be exacerbated by a range of constant maternal inflammatory and immune diseases many of which are known to be associated with enhanced levels of plasma CXCL12(Cruz-Orengo *et al*, 2011; Hansen *et al*, 2006; He *et al*, 2016; Ikawa *et al*, 2021). Further studies are required to understand the role of ACKR3 on pregnancy/fetus development in human, however, our findings provide new insights into the regulation of chemokines at the maternal-fetal interface and the importance of this regulation for development of the hematopoietic and immune systems in pups.

## Materials and methods

### Mice

WT C57BL/6J mice were from Charles River Laboratories. ACKR3-GFP reporter mice were from Jackson Labs (C57BL/6-Ackr3tm1Litt/J)(Cruz-Orengo *et al*., 2011). 7-10 week-old female mice were used for timed mating. Mice were maintained in specific pathogen-free facilities at the Beatson Institute and University of Glasgow. All experiments were performed under a UK Home Office Project License and approved by local ethical review committee at the University of Glasgow.

### *In vivo* experimental procedures

Ackr3^GFP/-^ females mated with genotype-matched males were injected with 0.5μg of AlexaFluor 647- conjugated human CXCL12. 1 or 3 hours after the injection placentas and embryonic blood were obtained for immunofluorescence staining and ELISA, respectively. For *in vivo* blockade of ACKR3, timed pregnant wild-type females were subcutaneously injected with either Captisol (Vehicle) or 30mg/kg of CCX771(Zabel *et al*., 2009) between E12.5 and E18.5. In addition, PBS or 0.5μg of recombinant mouse CXCL12 were intravenously injected into pregnant mice between E13.5 and E18.5. Pregnant females were weighed and monitored daily for parturition. For induction of an inflammatory response, 2-week-old offspring were intraperitoneally injected with PBS or 100μg of Deep Red *E.coli* Bioparticles. 24 hours after the injection peritoneal lavage and BM cells were obtained and analysed by flow cytometry. Pups were monitored daily and weighed weekly prior to the injection or tissue collection. Length of pups was defined as the distance from the tip of the nose to the start of tail.

### Digestion of placentas

For flow cytometry, placentas were separated into maternal decidua and placenta proper with fine forceps using a dissecting microscope. Tissues were incubated in 1ml of HBSS-based digestion cocktails containing 800μg/ml of dispase-II , 200μg/ml of collagenase-P, and 100μg/ml of DNase-I at 37°C at 1000rpm for 1 hour on a temperature-controlled shaker. 1ml of RPMI 1640 with 10% fetal calf serum (FCS) was added to digested tissues to neutralize enzymes and the tissues were filtered through 100μm cell strainers. After washing cells with PBS, cells were incubated with red blood cell lysis buffer and resuspended in FACS buffer (1% FCS and 2mM EDTA in PBS).

### Quantitative Real-time PCR

Total RNAs were isolated from maternal decidua and placentas using a PureLink RNA Mini Kit according to the manufacturer’s instructions. 100ng of RNAs were used for synthesis of cDNA using a High Capacity cDNA reverse transcription Kit(Dyer *et al*, 2019). qRT-PCR was performed using a PerfeCTa SYBR Green SuperMix on the ABI 7900HT. Data were normalized to the expression of *Gapdh*. Primer sequences are provided in supplementary information.

### Flow cytometry

Cell suspensions were incubated with Fc-block for 10 minutes at 4°C, and then stained with antibodies in FACS buffer for 30 minutes at 4°C. To define trophoblast cells, digested placental cells were stained with PerCP-Cy5.5-anti-mouse CD45, APC-anti-mouse CD9 and PE-anti-mouse CD71. To define HSC, BM and fetal liver (FL) cells were stained with APC-CD45, BV421-CD117 (c-Kit), PE-Cy7-Ly-6A/E (Sca-1) and lineage markers including biotin-anti-mouse CD3ε, B220, CD19, NK1.1, F4/80, CD11c and Ly-6G/Ly-6C (Gr-1). PerCP-Cy5.5-conjugated streptavidin was used for detection of biotinylated antibodies. To define monocytes, macrophages and neutrophils in peritoneal lavage and BM, cells were stained with anti-mouse BV785-CD11b, APC-Fire750-Ly6c, BV650- CD19, PE-Cy7-F4/80, BUV395-CD45 and BUV805-Ly6G. To define B cells, cells were stained with anti-mouse BV605-CD45, PerCP-Cy5.5-CD19, FITC-B220, BV421-IgD and APC-IgM. After staining with antibodies, cells were incubated with fixable viability dye eFluor780 or eFlour506 for 20 minutes at 4°C. Cells obtained were washed and then fixed with 2% paraformaldehyde (PFA). Flow data were obtained using a LSRII or Fortessa flow cytometer and analyzed by FlowJo software Ver.10.

### Immunofluorescence staining

Placentas were fixed with 4% PFA at 4°C overnight. The tissues were incubated with 30% sucrose for 2 hours at 4°C and then frozen in OCT. Frozen tissues were sectioned to 7μm with a cryostat. Sections were incubated with blocking buffer (1% BSA and 0.5% Tween-20 in PBS) for 15 minutes at RT, and then stained with biotin-anti-mouse CD9 or Rabbit anti-MCT4 at 4°C overnight. Sections were washed and incubated with Streptavidin-Alexafluor 594 or Alexafluor 594 anti-rabbit IgG. 40x- magnification Images were obtained using the Axioimager M2.

### Human samples

Placental samples were obtained at elective caesarean sections and placed in 10% neutral buffered formalin for fixation, processing and embedding. Written informed consent was obtained and the project was approved by the West of Scotland Research Ethics Committee (reference 17/WS/0174 and 19/WS/0111).

For immunohistochemistry, sections were deparaffined as above, then incubated with Bloxall to block endogenous peroxidase and heated in retrieval buffer (10mM citric acid buffer, pH6.0). The sections were incubated with 2.5% horse serum for 30 minutes at room temperature (RT) followed by staining with polyclonal anti-human ACKR3 antibody or isotype control at 4°C overnight. They were then incubated with horseradish peroxidase-conjugated secondary antibody for 30 minutes at RT and visualized using 3,3-diaminobenzibine for 30 seconds at RT. Sections were counterstained with hematoxylin followed by dehydration using ethanol and xylene. 40x magnification stitched Images were obtained using an EVOS FL auto2 microscope.

### Hematoxylin and eosin (H&E) staining

Placentas were fixed with 4% PFA at 4°C overnight. For H&E staining, the tissues were processed, embedded in paraffin and sectioned at 5μm. The sections were deparaffinised in xylene and rehydrated in serial ethanol gradients and then stained with hematoxylin and eosin. 20x magnification ‘stitched images’ were obtained using an EVOS FL auto2 microscope.

### ELISA

Embryonic blood was collected, following decapitation, into 1.5ml Eppendorf tubes filled with 20ml of 0.2M EDTA. The blood was centrifuged at 1,500xg for 10 minutes at 4°C to collect the plasma. The concentrations of CXCL12 were measured using mouse or human-specific CXCL12 Duoset ELISA kits according to manufacturer’s instructions. Total protein concentrations were measured using a Pierce BCA Protein Assay kit. The concentration of CXCL12 was normalized to the overall concentration of total proteins.

### Hematology study

Pups were euthanized with Dolethal at postnatal day 1 or 2. Blood was collected from jugular veins into EDTA-coated tubes filled with 20μl of 0.2M EDTA. Hematological analysis was performed by University of Glasgow Veterinary diagnostic services.

### Statistics

Two-tailed Student’s *t*-test, one-way ANOVA followed by Tukey’s test, and two-way ANOVA followed by Bonferroni’s test were performed using Prism9. *p* values of less than 0.05 were considered statistically significant. All data are presented as mean±SD.

### Data sharing statement

For original data, please contact

## Acknowledgements

This work was supported by grants to GJG from the Wellcome Trust (217093/Z/19/Z) and the MRC (MR/V010972/1). We thank all staff working in animal facilities, flow cytometry for technical assistance and Diagnosis service in University of Glasgow. We thank Prof. Katrin Ottersbach for technical advice. We thank all our colleagues in the chemokine research group in University of Glasgow and at ChemoCentryx particularly Tom Schall and Jim Campbell.

## Author contribution

G.J.G. and A.F. designed the study, analyzed data, and wrote the manuscript. A.F, G.J.W., E.P., L.K.F. and M.P. performed experiments and analyzed data. H.J. obtained human placentas and sectioned samples. G.J.G. supervised the study. All authors discussed the results and reviewed and revised the manuscript.

